# Early *Hox* Gene Expression in Echinoderms

**DOI:** 10.1101/2024.08.26.609757

**Authors:** Olga V. Ezhova, Natalya V. Ageenko, Konstantin V. Kiselev, Anastasiya I. Lukinykh, Vladimir V. Malakhov

## Abstract

Breakages of sequential *Hox* expression in Echinodermata may reflect evolutionary changes in the body plan. We quantified *Hox* gene expression at early larval stages, i.e., blastula at 13 hpf, gastrula at 35 hpf, prism stage at 46 hpf, and echinoplutei at 4 and 9 dpf, in the sea urchin with indirect development, *Strongylocentrotus intermedius*. The design of specific primers for real-time PCR (q-RT–PCR) for *S. intermedius* was performed. At blastula stage, only *SiHox7* (medial *Hox* group), *SiHox11/13b* and *SiHox11/13c* (posterior *Hox* group) were expressed. At gastrula and prism stages, the expression of *SiHox* genes was lower than that in unfertilized eggs. Significant increase in the expression of all *SiHox* genes was observed in 4-day pluteus with further increase of *SiHox1, SiHox8, SiHox9/10*, and *SiHox11/13b* expression in 9-day pluteus.

In other echinoderms, the genes of anterior *Hox* group are nearly not expressed in the early stages of development. At blastula, gastrula and prism stages, the genes of medial *Hox* group (mainly *Hox7* and *Hox8*) and posterior *Hox* group (mainly *Hox11/13a, 11/13b, 11/13c*) are significantly expressed. This breakage of sequential *Hox* expression in echinoderm ancestors may be caused by loss of metameric gill slits, which are the most important synapomorphy of deuterostomes.

## Introduction

In Eumetazoa, the temporal and spatial collinearity of *Hox* gene expression reflects the sequential formation of different body structures along the anteroposterior axis of an animal (Akam 1989; Arendt 2018). In addition, the posterior prevalence is described for the activity of Hox proteins when the more 5’, or posterior, proteins counteract the function of the more 3’, or anterior, proteins (Duboule 2007). In this regard, the relationship between the breakage of collinearity and the pattern of an animal’s body plan is an interesting challenge. Among the different phyla, the groups with complicated, intricate body plans demonstrated deviations from the stepwise order of *Hox* expression. In molluscs, the clades Bivalvia, Gastropoda and Cephalopoda have undergone significant morphological changes in the ancestral body plan, and their representatives have lost collinear *Hox* expression (Wollesen et al. 2018). A lack of temporally collinear *Hox* expression is observed in some phoronids and brachiopods (Schiemann et al. 2017; Gąsiorowski and Hejnol 2020). In deuterostomes, tunicates are known for their curious structure and for their dispersed *Hox* cluster organization accompanied by retained spatially coordinated expression with loss of temporal expression (Ikuta et al. 2004; Seo et al. 2004).

Echinoderms have been characterized by Edward E. Ruppert and coauthors as “variations on celestial stars that fell to Earth as extraterrestrials, so extraordinary is their form and function” (Ruppert et al. 2004, P. 873). Vary breakages of sequential expression of *Hox* genes have been described for echinoderm development. According to current data, in the echinoderm blastula stage, no expression of the anterior *Hox* gene group is observed, and only two genes of the posterior *Hox* group are expressed in the holothuroid *Apostichopus japonicus* (see Kikuchi et al. 2015), and only one gene of the medial *Hox* group and one or two genes of the posterior *Hox* group are expressed in the echinoids *Strongylocentrotus purpuratus* and *Peronella japonica* (see Arenas-Mena et al. 1998; Tsuchimoto and Yamaguchi 2014). In the crinoid *Metacrinus rotundus, Hox* expression is not detected in the blasula stage (Hara et al. 2006). In the echinoderm gastrula stage, *Hox* expression is still weak and increases during later developmental stages. This breakage in temporal collinearity may be related to maximal indirect development, which is an ancestral pattern for echinoderms (Arenas-Mena et al. 1998), or to the unusual gene order in the echinoderm *Hox* cluster (Cameron et al. 2006) (Fig. 1). In both cases, it may also reflect the strong morphological changes that the echinoderm body plan has undergone during the evolution. The reported data on *Hox* expression in the early developmental stages of echinoderms are limited by several species and accumulate slowly, being continually corrected and forming the basis of new evolutionary hypotheses.

**Figure 1.**
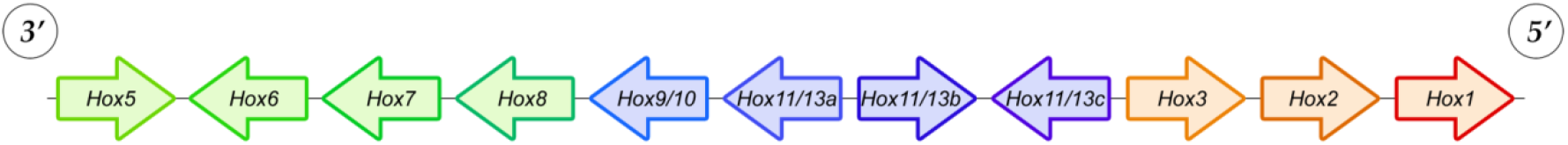
The *Hox* gene cluster of the sea urchin *Strongylocentrotus purpuratus* with a unique gene order in which the paralog groups are not expressed in a sequential manner (Cameron et al. 2006). The colors indicate the *Hox* gene groups: red – anterior, green – central, blue – posterior.

Here, we provide our results of quantify *Hox* cluster gene expression at the early larval stages, from blastula to 9-day echinopluteus, in the sea urchin with indirect development, *Strongylocentrotus intermedius* (Fig. 2A, B). The developmental stages used for the experiment were unfertilized eggs, blastulae at 13 hours post fertilization (hpf), gastrulae at 35 hpf, prism larvae at 46 hpf, and plutei at 4 days pf (dpf) and 9 dpf (Fig. 2C). The *Hox* cluster of *Strongylocentrotus* sea urchins contains 11 genes with the order 5’-*Hox1, 2, 3, 11/13c, 11/13b, 11/13a, 9/10, 8, 7, 6, 5*-3’, where the anterior genes (*Hox1, 2*, and *3*) lie nearest the posterior genes (*Hox9/10, 11/13a, 11/13b*, and *11/13c*) (Fig. 1) (Cameron et al. 2006). The *Hox* gene expression was quantified individually for each gene in relative units (r.u.), taking the level of amplification of the gene in unfertilized eggs as the conditional value ‘1’. In addition, we compared early *Hox* gene expression in *S. intermedius* with that in other studied echinoderm species, which allowed us to suggest the evolutionary significance of the temporal *Hox* expression order in Echinodermata.

**Figure 2.**
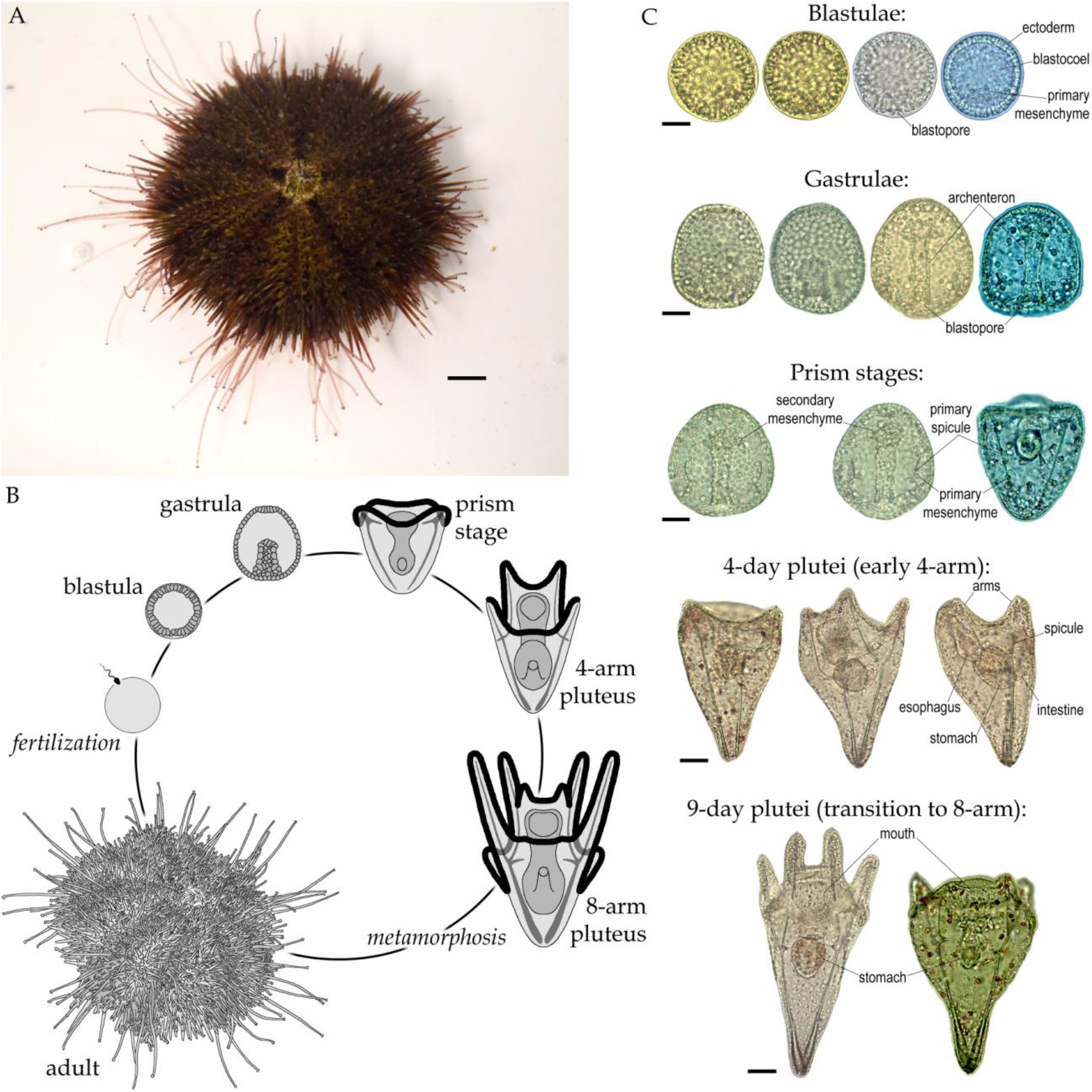
Sea urchin *Strongylocentrotus intermedius* (**A**), its life cycle (**B**) and developmental stages, which were used in this work (**C**). Scale bars, 1 cm (**A**), 20 μm (**C**).

## Results

We quantified the expression of all 11 *Hox* genes in the early stages of development of *S. intermedius* (Fig. 3).

**Figure 3.**
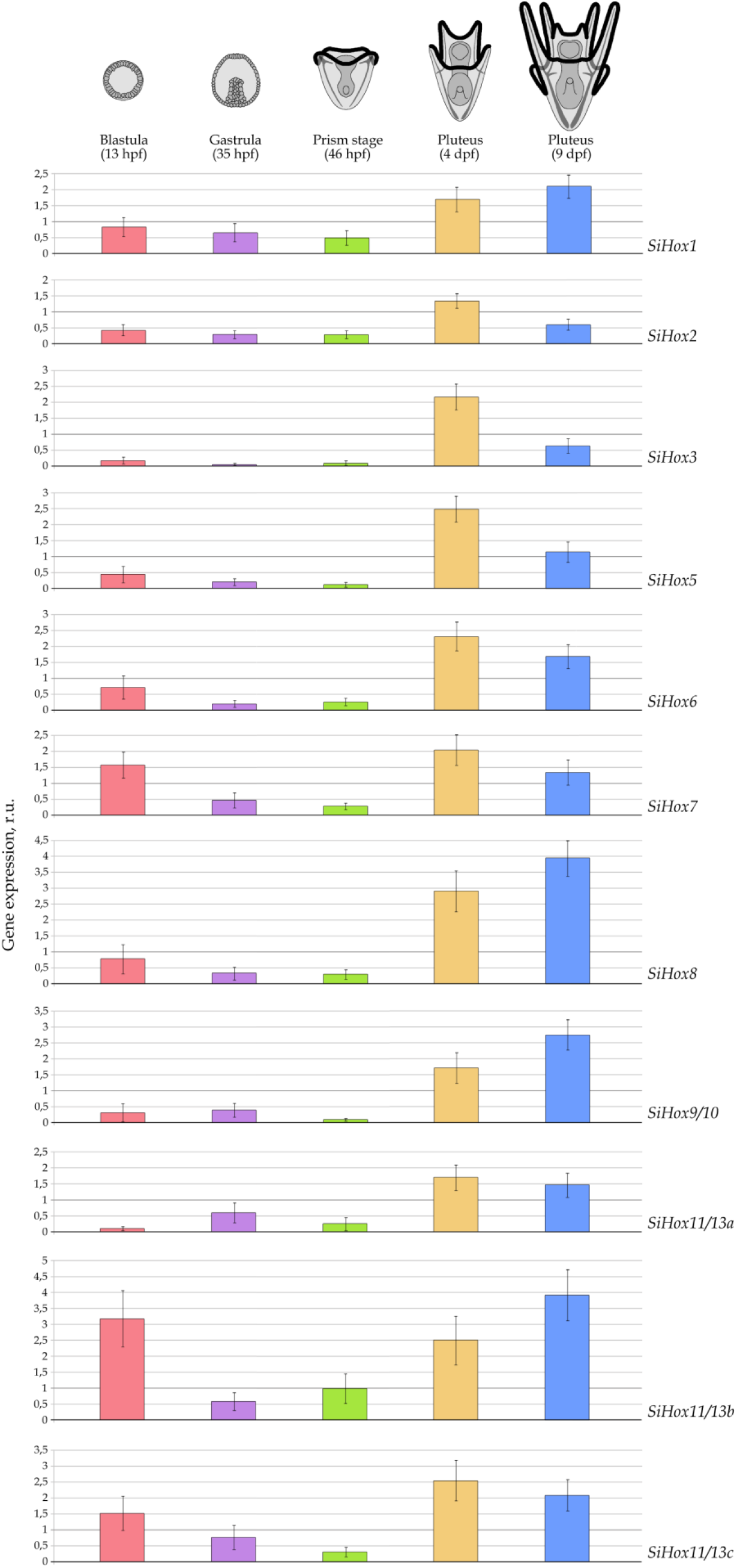
Quantify expression of each gene of the *Hox* cluster at the early stages of larval development of *Strongylocentrotus intermedius* (in relative units, r.u., where the level of amplification of the gene in unfertilized eggs is taken as the conditional value ‘1’).

The expression of *SiHox1, SiHox8*, and *SiHox9/10* was lower than that in unfertilized eggs at the blastula, gastrula and prism stages. A significant increase in the expression of these genes was detected in the 4-day pluteus, with a further increase in the 9-day pluteus. The expression of *SiHox2* and *SiHox3* exceeded that of unfertilized eggs only in the 4-day pluteus. The expression of *SiHox5, SiHox6*, and *SiHox11/13a* was greater than ‘1’ in the 4-day pluteus and 9-day pluteus, with a slight decrease in the latter.

Early expression exceeding the level of unfertilized eggs was observed for *SiHox7, SiHox11/13b*, and *SiHox11/13c* at the blastula stage. The expression of all three genes decreased to levels lower than ‘1’ at the gastrula and prism stages and again exceeded the ‘1’ level in 4-day pluteus; however, for *SiHox11/13b*, the expression at this stage did not exceed the blastula level. In 9-day pluteus, the expression of *SiHox7* and *SiHox11/13c* decreased slightly, remaining greater than ‘1’, and the expression of *SiHox11/13b* continued to increase.

The general pattern of early *SiHox* expression was as follows. A decrease in expression at the blastula stage in comparison with that in unfertilized eggs was observed for *SiHox* genes, except for *SiHox7* (medial *Hox* group), *SiHox11/13b* and *SiHox11/13c* (posterior *Hox* group); the expression of these three genes conversely increased. At the gastrula and prism stages, the expression of *SiHox* genes continued to decrease, but a slight increase in expression was observed for *SiHox11/13a* at the gastrula stage and for *SiHox11/13b* at the prism stage. Significant intensification of the expression of all the *SiHox* genes was observed in the 4-day pluteus. In 9-day pluteus, the expression of *SiHox1, SiHox8, SiHox9/10*, and *SiHox11/13b* continued to increase, whereas the expression of other *SiHox* genes slightly decreased (Fig. 3).

## Discussion

Early expression of the medial gene *Hox7* and the posterior genes *Hox11/13b* and *Hox11/13c* at the blastula stage is typical for the development of other sea urchins (Fig. 4). In ‘regular’ sea urchin with indirect development, *S. purpuratus*, a high level of expression at the blastula stage was reported for two genes, *SpHox7* (24 hpf) and *SpHox11/13b* (12 hpf and 24 hpf) (Angerer et al. 1989; Dobias et al. 1996; Arenas-Mena et al. 1998); the expression of *SpHox 11/13c* has not been analysed in the cited works. In a sand dollar with direct development, *Peronella japonica*, early activation at the blastula stage (6–10.5 hpf) was described for three genes: *PjHox7, PjHox11/13b* and *PjHox11/13c* (Tsuchimoto and Yamaguchi 2014). These three genes are expressed at the blastula stage (13 hpf) in *S. intermedius*. In the holothuroid *Apostichopus japonicus*, the genes *AjHox11/13a* and *AjHox11/13b* are expressed at the blastula stage (12hpf); in hatching embryos (20 hpf) of *A. japonicus*, the expression of two more genes, *AjHox7* and *AjHox8*, is activated (Kikuchi et al. 2015).

**Figure 4.**
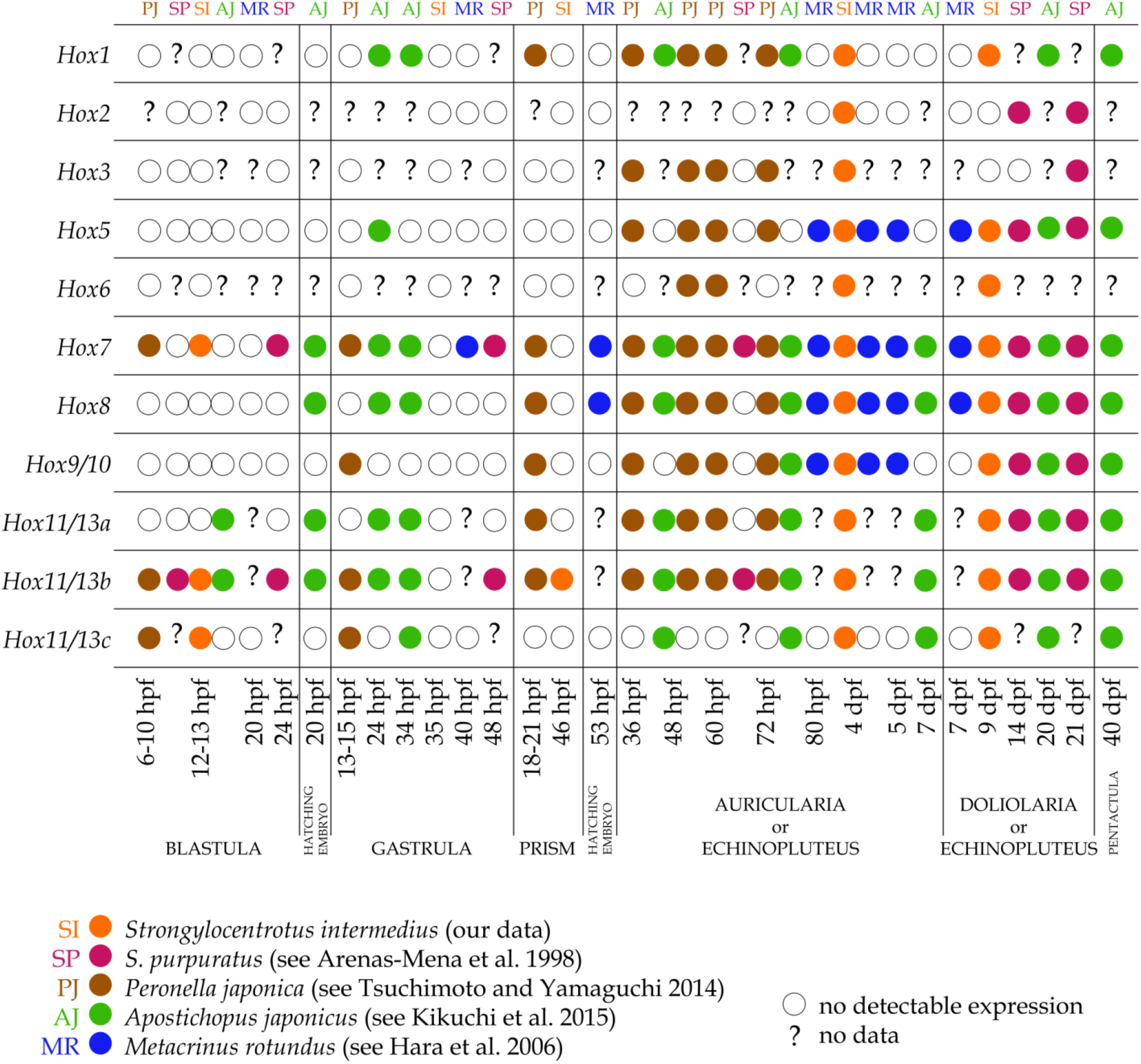
Expression of *Hox* genes during Echinodermata development.

At the gastrula stage in *S. purpuratus* (48 hpf), the expression of only *SpHox7* and *SpHox11/13b* continues, although the latter is greatly decreased (Arenas-Mena et al. 1998). In *P. japonica, PjHox7, PjHox11/13b* and *PjHox11/13c* expression continues, and *PjHox9/10* expression is added (Tsuchimoto and Yamaguchi 2014). However, in *S. intermedius*, the expression of *Hox* genes is not detected at the gastrula stage, and even *SiHox7, SiHox11/13b*, and *SiHox11/13c* expression is lower than that in unfertilized eggs (Fig. 3). In *A. japonicus*, all genes activated at the blastula stage continue to be expressed at the gastrula stage (24 and 34 hpf), and in addition, slight *AjHox1, AjHox5*, and *AjHox11/13c* expression is detected (Kikuchi et al. 2015). In the crinoid *Metacrinus rotundus, Hox* expression was first detected at the gastrula stage (40 hpf), *MrHox7. MrHox8* expression, in addition to *MrHox7* expression, is detected in hatching embryos of *M. rotundus* (2.2 dpf) (Hara et al. 2006) (Fig. 4).

*Hox* gene expression at the prism stage has been described only for *P. japonica* (18 hpf and 21 hpf). *PjHox1, PjHox8*, and *PjHox11/13a* expression was first detected at this stage, and *PjHox11/13c* became undetectable (Tsuchimoto and Yamaguchi 2014). In *S. intermedius*, only *SiHox11/13b* expression reached the level of unfertilized eggs at the prism stage (Fig. 3).

In summary, the genes of the anterior *Hox* group are nearly not expressed in the early stages of Echinodermata development (Fig. 4). At the blastula, gastrula and prism stages, the genes of the medial *Hox* group (mainly *Hox7* and *Hox8*) and posterior *Hox* group (mainly *Hox11/13a, 11/13b*, and *11/13c*) were significantly expressed. In Echinoidea, the genes of the anterior *Hox* group and *Hox5* and *Hox6* of the medial *Hox* group begin to be expressed only at the pluteus stage (Fig. 4). For the holothuroid *A. japonicus, Hox2, Hox3*, and *Hox6* expression has not been studied; slight *AjHox1* and *AjHox5* expression was first detected at the gastrula stage; however, a significant increase in the expression of these two genes was detected only at the doliolaria stage (Kikuchi et al. 2015). In the crinoid *M. rotundus, MrHox1, MrHox2, MrHox4*, and *MrHox11/13c* expression was not detected; the expression of *Hox3, Hox6, Hox11/13a*, and *Hox11/13b* was not studied. *MrHox5* and *MrHox9/10* expression was first detected at the preauricularia stage (3 dpf), in addition to *MrHox7* and *MrHox8* expression (Hara et al. 2006).

This ‘silencing’ of the anterior *Hox* genes in Echinodermata may reflect the complex evolutionary transformations of the echinoderm body. Echinodermata together with Hemichordata belong to the clade Ambulacraria, which is a sister to the phylum Chordata within the group Deuterostomia (Fig. 5A). The most important synapomorphy of deuterostomes is the metameric pharyngeal gill slits and corresponding external gill pores (Ezhova and Malakhov 2022). The gill slits and gill pores are present in chordates and hemichordates, but this branchial apparatus is lost by modern echinoderms (Fig. 5A). It is known that in triploblastic Bilateria, the *Hox* gene system regulates the metameric organization of the trunk, whereas the preoral and perioral (=tentacular) regions are free of *Hox* gene expression (Finkelstein and Boncinelli 1994; Irvine and Martindale 2000; Kuratani 2005; Lowe et al. 2003; Olsson et al. 2005; Aronowicz and Lowe 2006; Kulakova et al. 2007; Fröbius et al. 2008; Bakalenko et al. 2013; Janssen et al. 2014; Gonzales et al. 2017; Andrikou et al. 2019; Gąsiorowski and Hejnol 2020; etc.). In Chordata and Hemichordata, metameric pharyngeal gill slits are the main markers of the anterior trunk region (Fig. 5A). It may be suggested that *Hox* expression relates to the formation of branchial apparatus in deuterostomes. In chordates, the expression of anterior *Hox* genes is involved in the patterning of the pharyngeal arches (Hunt et al. 1991; Wilkinson 1993); the effect of excess retinoic acid on the expression of anterior *Hox* genes causes pharyngeal arch abnormalities and the failure of mouth and gill slits to form (Morriss-Kay et al. 1991; Holland and Holland 1996).

**Figure 5.**
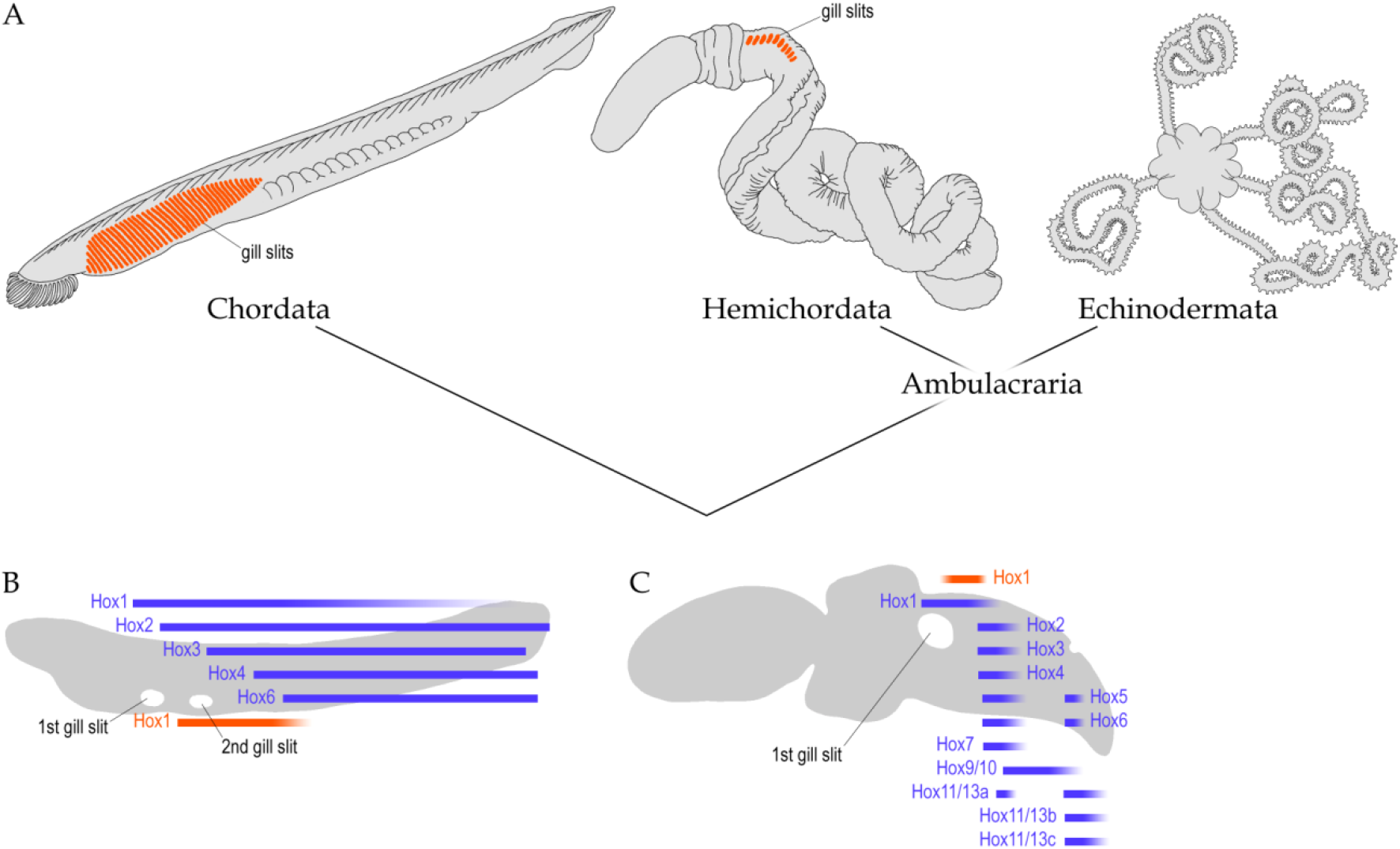
Presence of the metameric pharyngeal gill slits in modern Deuterostomia (**A**) and *Hox* gene expression (blue – in the CNS, based on Schubert et al. 2006; orange – in the endoderm, based on Schubert et al. 2005) in cephalochordates (**B**) and in hemichordates (blue – in the basiepithelial nerve net; orange – in the endoderm, based on Aronowicz and Lowe 2006) (**C**) in relation to the first gill pores.

In the developing endoderm of Cephalochordata, the anterior limit of *Hox1* expression is where the first gill slit will form (Schubert et al. 2005). Moreover, in cephalochordates, the *Hox* genes are collinearly expressed according to the position of the somites (Fig. 5B) (Schubert et al. 2006). In hemichordates, expression of *Hox1* in the endoderm is detected just posterior to the first gill slit (Fig. 5C) (Aronowicz and Lowe 2006). However, echinoderm ancestors have lost the pharyngeal gill apparatus and it could cause the breakage of sequential *Hox* expression, in which the anterior *Hox* genes are not detected at early developmental stages.

## Materials and Methods

The sea urchins *S. intermedius* were collected in the East Bay of the Sea of Japan. The animals were kept in baths with running aerated water, and before starting the experiment, they were washed two to three times with filtered seawater treated with ultraviolet light. Spawning was induced by injecting 1–2 ml of 0.5M potassium chloride solution into the cavity of the Aristotelian lantern. The larvae were obtained by artificial insemination and further cultured at 18ºC. All experiments on the fertilization and development of sea urchins were performed at the Marine Biological Station «Vostok» of the A.V. Zhirmunsky National Scientific Center for Marine Biology of the Far Eastern Branch of the Russian Academy of Sciences (NSCMB FEB RAS).

Total RNA (tRNA) was isolated from the obtained eggs and larvae of the necessary stages of *S. intermedius* according to the methods described in detail in Kiselev et al. (2013). To the sediment of the collected eggs and larvae (blastula, gastrula, prism and pluteus) of *S. intermedius*, 1.4 ml of buffer for tRNA isolation was added, and the mixture was thoroughly mixed. The resulting mixture was incubated in a thermostat (+65ºC) for 5 min, after which a chloroform solution (500 μl) was added, mixed and centrifuged for 10 min (+4ºC, 10000 rpm). Next, the supernatant was selected and transferred to a new test tube with a solution of lithium chloride (250 μl). The resulting mixture was incubated for 1 day (the incubation time can be increased to 3 days) at +4ºC (refrigerator), after which it was centrifuged for 15 min (+4ºC, 10000 rpm). The supernatant was drained, and 100 μl of water was added to the precipitate, which was then thoroughly mixed. Next, 2.5–3 volumes of 95% ethyl alcohol (300 μl) were added to the resulting solution and the mixture was kept at –20ºC (refrigerator) for 20–30 min (1 day is allowed). After centrifugation (15 min, +4ºC, 10000 rpm), the supernatant was removed, and 500 μl of 70% ethyl alcohol solution was added to the precipitate; after thorough mixing, the alcohol was removed with a pipette tip. The resulting precipitate was dried in a thermostat (30 min, +37ºC) and 120 μl of water was added to it. The quantitative assessment of the isolated tRNA was carried out spectrophotometrically (Shimadzu, Japan).

Complementary DNA (cDNA) was obtained via a reverse transcription kit (Silex, Moscow, Russia) with 1–3 μg of isolated tRNA. Reverse transcription polymerase chain reaction (RT–PCR) requires 50 μl of a reaction mixture containing one RT buffer (RT–PCR buffer), 0.24 mm each of 4 deoxynucleotide triphosphates (dNTP), 0.2 μm oligo-(dT)15 primer, and 200 units of MMLV reverse transcriptase actions. The reaction was carried out at 37ºC for 2 hours. Samples of the obtained products (0.5 μl) were then amplified via PCR. To normalize the differences in the quality and quantity of the obtained cDNA used in each real-time PCR experiment (q-RT–PCR), the actin gene (GenBank: DQ229162) and the ubiquitin gene (GenBank: LOC754856) were used as endogenous controls for the *S. intermedius* sea urchin. The data were obtained from five independent experiments. The primers used for the actin and ubiquitin genes for q-RT–PCR were previously described (Ageenko et al. 2011).

Sequencing was carried out on an ABI 3130 Genetic Analyser sequencer (Applied Biosystems, USA) at the Federal Scientific Center of East Asia Terrestrial Biodiversity, Far Eastern Branch of the Russian Academy of Sciences. Each transcript was sequenced at least three times.

For the theoretical design of specific primers for q-RT–PCR genes of the *S. intermedius* sea urchin, we analysed the nucleotide sequence of genes of the *Hox* cluster of the closely related species *S. purpuratus* (see Tu et al. 2012, 2014). All the data and related input and visualization tools are available through SpBase, a publicly available database of the sea urchin genome (http://www.spbase.org/SpBase/rnaseq/), and via the NCBI Bioproject database (http://www.ncbi.nlm.nih.gov/bioproject), registration number PRJNA81157. On the basis of the obtained sequencing data, the design of specific primers for q-RT–PCR was performed (Table 1). The NCBI Primer Blast program was used to create primers.

**Table 1.**
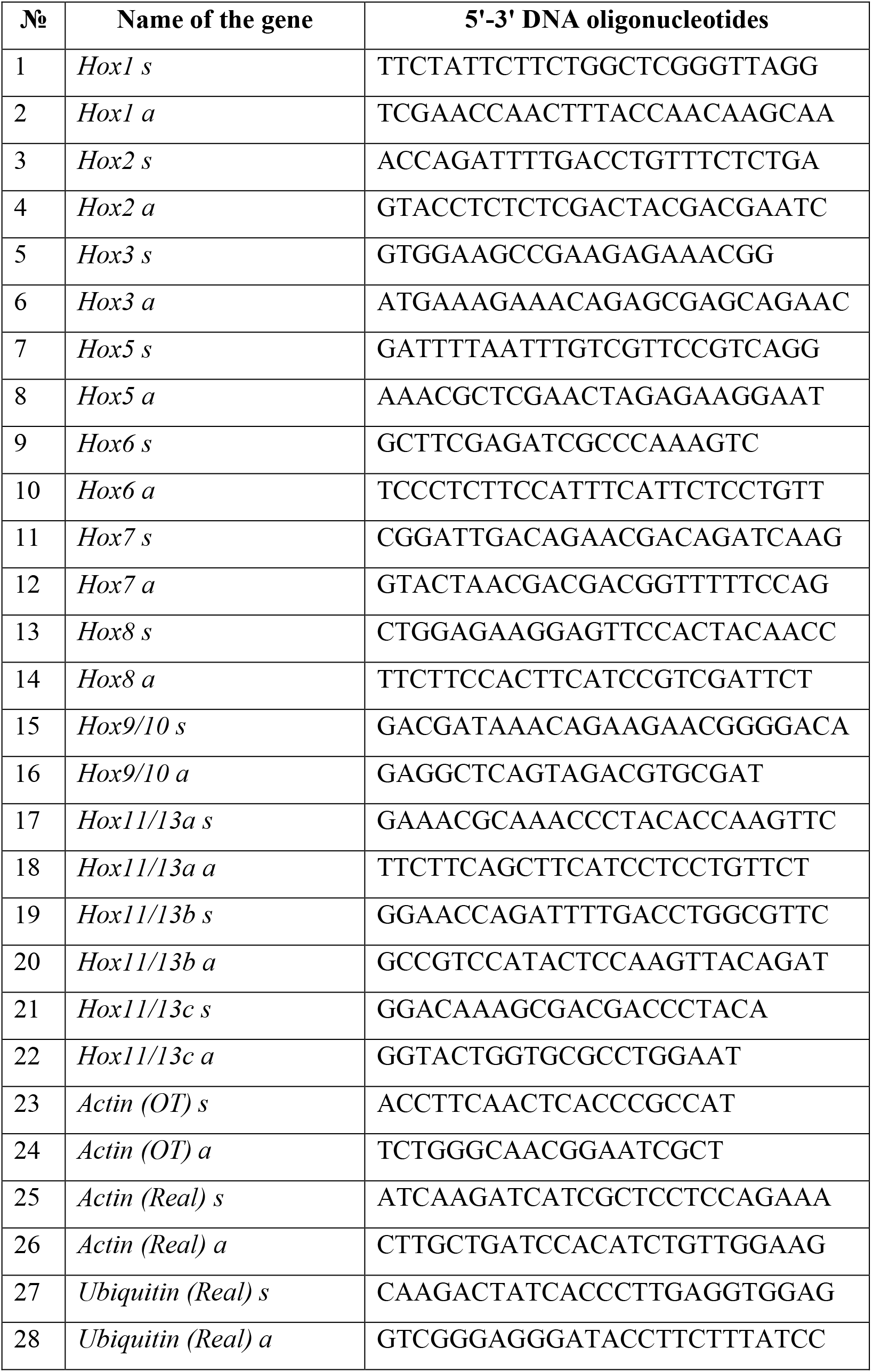
List of primers (5′-3′) used to amplify the *Hox* genes, the *Actin* gene in OT–PCR, the *Actin* gene and the *Ubiquitin* gene in q-RT–PCR analysis.

## Competing Interest Statement

The authors declare that they have no conflict of interest or competing financial interests.

All the authors have approved the current version of the manuscript and have approved its submission to *Genes & Development*.

## Ethics approval

All applicable international, national, and/or institutional guidelines for the care and use of animals were followed. In accordance with Directive 2010/63/EU of 22 September 2010 on the protection of animals used for scientific purposes, Chapter 1, Paragraph 3, the requirements of bioethics do not apply to the object of this study.

## Acknowledgments

The authors are grateful to the staff of the diving service of NSCMB FEB RAS for collecting specimens of the sea urchin *S. intermedius* for the study. We thank Anatoliy Drozdov (NSCMB FEB RAS, Vladivostok, Russia) for valuable consultation on the development of sea urchins. We thank Vladimira Shuklova and Alarikh Shuklov for providing an opportunity to prepare the manuscript.

The study was supported by the Russian Science Foundation (RSF), project no. 23-14-00047.

## Author Contributions

Ezhova O.V. – drafting the work, manuscript preparation, figures preparation, analysis and interpretation of the obtained data, references search and analysis;

Ageenko N.V. – material collection and photographing, tRNA isolation, cDNA obtaining, sequencing, design of specific primers, conducting PCR and q-RT–PCR analysis, manuscript improvement;

Kiselev K.V. – tRNA isolation, cDNA obtaining, sequencing, design of specific primers, conducting PCR and q-RT–PCR analysis, manuscript improvement;

Lukinykh A.I. – material collection and photographing, manuscript improvement;

Malakhov V.V. – conception and design of the work, references search and analysis, manuscript improvement.

